# Unveiling genome plasticity and a novel phage in *Mycoplasma felis*: genomic investigations of four feline isolates

**DOI:** 10.1101/2023.12.16.572022

**Authors:** Sara M. Klose, Alistair R. Legione, Rhys N. Bushell, Glenn F. Browning, Paola K. Vaz

## Abstract

*Mycoplasma felis* has been isolated from diseased cats and horses, but to date only a single fully assembled genome of this species, of an isolate from a horse, has been characterised. This study aimed to characterise and compare the completely assembled genomes of four clinical isolates of *M. felis* from three domestic cats, assembled with the aid of short and long read sequencing methods. The completed genomes encoded a median of 759 open reading frames (min, 743, max 777) and had a median average nucleotide identity (ANI) of 98.2% with the genome of the available equid origin reference strain. Comparative genomic analysis revealed the occurrence of multiple horizontal gene transfer (HGT) events and significant genome reassortment. This had resulted in the acquisition or loss of numerous genes within the Australian felid isolate genomes, encoding putative proteins involved in DNA transfer, metabolism, DNA replication, host cell interaction, and restriction modification systems. Additionally, a novel mycoplasma phage was detected in one Australian felid *M. felis* isolate by genomic analysis and visualised using cryo-transmission electron microscopy. This study has highlighted the complex genomic dynamics in different host environments. Furthermore, the sequences obtained in this work will enable the development of new diagnostic tools, and identification of future infection control and treatment options for the respiratory disease complex in cats.

**Data summary:** All genome data for this study have been deposited in GenBank under BioProject PRJNA906261. Genome assemblies, as well as Illumina and Oxford Nanopore sequence reads for each isolate, can be found under their respective BioSamples: SAMN32182834 (isolate 047), [SAMN32182835](https://dataview.ncbi.nlm.nih.gov/object/SAMN32182834) (isolate 219), [SAMN32182836](https://dataview.ncbi.nlm.nih.gov/object/SAMN32182834) (isolate 329 and associated phage), and [SAMN32182837](https://dataview.ncbi.nlm.nih.gov/object/SAMN32182834) (isolate 632). The authors confirm all supporting data and protocols have been provided within the article.

**Impact statement:** *Mycoplasma felis* is commonly associated with clinical cases of conjunctivitis and feline respiratory disease complex in cats, the leading cause of euthanasia in animal shelters. In the absence of vaccines, infection control is currently limited to the prolonged treatment with antimicrobials. Prior to this study there was only one complete genome assembly of an isolate of *M. felis*, which had been obtained from a horse. This study has provided the first high quality hybrid assembled genomes of *M. felis* isolates from cats. This work adds four new genomes from clinical cases, as well as the identification and validation of the presence of a novel phage that utilises the mycoplasma translation code. The genomic data presented here can assist future projects investigating improved diagnostics and development of new treatment options for this significant feline pathogen.

## Introduction

Mycoplasmas are the smallest free-living bacteria and have some of the smallest bacterial genomes (0.58 – 1.38 Mb). They are thought to approach the minimal gene sets essential for independent life, making them ideal model organisms for studies on the fundamental requirements of a living cell [1]. They evolved from Gram positive bacterial ancestors 600 million years ago [2] and have undergone considerable genome reduction during their evolution [3, 4]. They lack a cell wall, are pleomorphic and can infect and cause acute and chronic disease in a diverse range of animal and plant species [5].

Some mycoplasmas have been found to exhibit significant genetic plasticity and to evolve rapidly, mainly driven by intra-species spontaneous mutation and recombination or horizontal gene transfer (HGT) between and within species, facilitating swift adaptation to environmental changes, enhancing survival and/or virulence [6–9]. Studies examining the genomic stability of *Mycoplasma synoviae* have revealed that mutations occur frequently during infection in birds [10], and that rapid thermoadaptive evolution can occur *in vitro* and *in vivo* [11], presumably to regain fitness and pathogenicity [12, 13]. HGT appears to occur more frequently between species that share an ecological niche, irrespective of their phylogenetic divergence. Examples of significant HGT include the transfer of the genes encoding the phase variable VlhA cell surface lipoproteins between the poultry pathogens *Mycoplasma gallisepticum* and *M. synoviae* [14], and the transfer of several cell surface lipoproteins involved in host colonisation between the ruminant pathogens *Mycoplasma capricolum* subsp. *capricolum* and *Mycoplasma mycoides* subsp. *Mycoides*, and *Mycoplasma agalactiae* and *Mycoplasma bovis* [15].

At least four species of mycoplasmas have been isolated from cats (*Mycoplasma felis*, *Mycoplasma gateae*, *Mycoplasma arginini*, and *Mycoplasma feliminutum*), in most cases as commensals. *M. felis* in particular is primarily an opportunistic pathogen, able to infect and be isolated from the upper respiratory tract. It has been associated with conjunctivitis and, increasingly, feline respiratory disease [16–19]. Less commonly, *M. felis* has also been associated with respiratory disease in horses [20] although it is unclear how bacterial infection mechanisms and pathogenesis may vary between hosts as infection by *M. felis* is under-explored in both cats and horses. Feline respiratory disease complex (FRDC) is the leading cause of euthanasia of cats in animal shelters [21] and in addition to *M. felis* is also commonly associated with coinfections with several viral (feline herpesvirus 1 and feline calicivirus) and bacterial pathogens (*Chlamydia felis*) [22, 23]. *M. felis* infections are common and difficult to eradicate in most cat populations, and the ability of this organism to persist in an animal, likely due to intracellular invasion, enhances opportunities for recurrent opportunistic infections. Vaccines against *M. felis* are not currently available and treatment of bacterial infection is limited to prolonged treatment of up to eight weeks with tetracyclines or fluoroquinolones, which is arguably a risk for antimicrobial resistance (AMR) development [24, 25]. Currently, full genomes of three isolates of this species have been published: strain Myco-2 (GenBank accession number: AP022325.1), an isolate from a case of respiratory disease in a horse in Japan [26], which has been completely assembled, and multiple assemblies of isolates 16-040612-4 (CP104192, CP104193, CP104194, CP11027) and 16-057065-4 (CP103988, CP103993, CP110269), which were each isolated from feline bronchoalveolar lavages in Canada [27]. Additionally, there is an incomplete genome of ATCC strain 23391, assembled into scaffolds (GenBank Biosample: SAMN02841218).

The aim of this study was to sequence and assemble four high quality, complete *M. felis* genomes, using hybrid assembly methods. These genomes represent the first complete assemblies of feline isolates of this species. The objective was to significantly expand on our limited understanding of the genomic structure and plasticity of *M. felis*, which may lead to the development of improved diagnostic tests and vaccines, as has been seen in other mycoplasma species.

## Methods

### Mycoplasma isolation

Swabs were collected for bacteriological culture from clinical cases of respiratory disease (3 cases) and an unresolved post-surgical infection (1 case) (Table 1). Swabs were used to inoculate sheep blood agar (SBA) plates, which were incubated aerobically at 37°C. Following observation and presumptive identification of small mycoplasma-like colonies, single colonies were transferred to Mycoplasma Broth (MB) containing 10% swine serum (Sigma-Australia) and 0.01% nicotinamide adenine dinucleotide (NAD) (Sigma-Australia), based on the formulation of Frey’s medium, with minor modifications [28]. Acidification of the MB medium was used to confirm the growth of *Mycoplasma* spp. and the cultures were stored at -80°C for future investigation.

**Table 1.**
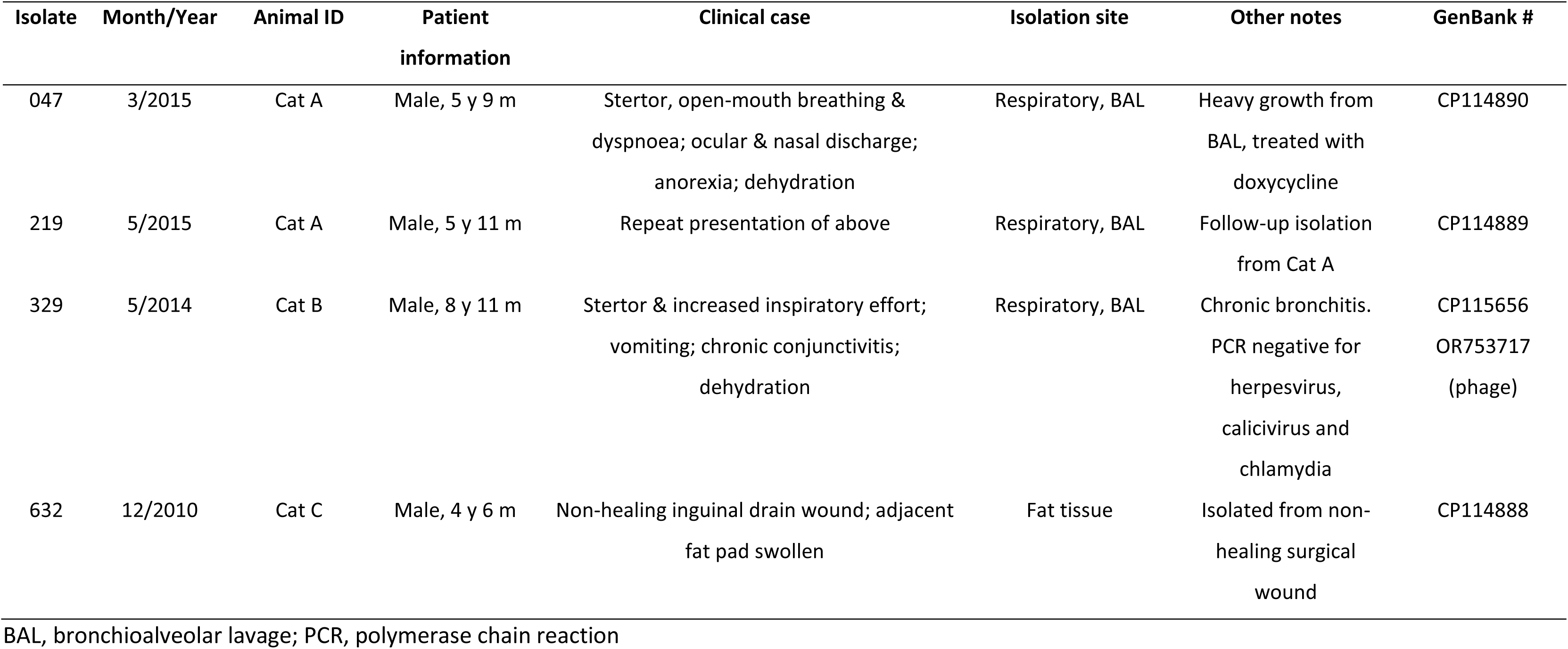
*Mycoplasma felis* isolates sequenced in this study.

| Isolate | Month/Year | Animal ID | Patient<br>information | Clinical case | Isolation site | Other notes | GenBank # |
| --- | --- | --- | --- | --- | --- | --- | --- |
| 047 | 3/2015 | Cat A | Male, 5 y 9 m | Stertor, open-mouth breathing &<br>dyspnoea; ocular & nasal discharge;<br>anorexia; dehydration | Respiratory, BAL | Heavy growth from<br>BAL, treated with<br>doxycycline | CP114890 |
| 219 | 5/2015 | Cat A | Male, 5 y 11 m | Repeat presentation of above | Respiratory, BAL | Follow-up isolation<br>from Cat A | CP114889 |
| 329 | 5/2014 | Cat B | Male, 8 y 11 m | Stertor & increased inspiratory effort;<br>vomiting; chronic conjunctivitis;<br>dehydration | Respiratory, BAL | Chronic bronchitis.<br>PCR negative for<br>herpesvirus,<br>calicivirus and<br>chlamydia | CP115656<br>OR753717<br>(phage) |
| 632 | 12/2010 | Cat C | Male, 4 y 6 m | Non-healing inguinal drain wound; adjacent<br>fat pad swollen | Fat tissue | Isolated from non-healing surgical<br>wound | CP114888 |

BAL, bronchioalveolar lavage; PCR, polymerase chain reaction

### DNA extraction, sequencing, and genome assembly

Mycoplasma isolates were cultured axenically in MB for 18 h at 37°C prior to DNA extraction. Bacteria were pelleted and the medium removed, and then resuspended in the appropriate extraction kit lysis buffer. The DNA for short read sequencing was extracted using the DNeasy Blood and Tissue Kit (QIAGEN), following the manufacturer’s protocol, and 100 ng of the extracted DNA was used to prepare sequencing libraries using Illumina’s Nextera Flex DNA Library Prep Kit. Sequencing was performed on the NovaSeq platform at Charles River Laboratories, Victoria, Australia, to generate 150 bp paired-end reads. Two rounds of Oxford Nanopore Technologies (ONT) long read sequencing were performed. The DNA was extracted using the Promega Wizard DNA Purification Kit (Round 1) or the Promega Wizard High Molecular Weight DNA Purification Kit (Round 2). DNA libraries were generated using the ONT Rapid Barcoding (SQK-RBK004) Kit. Sequencing was performed on a MinION Mk1b device fitted with a FLO-MIN106D Flow Cell (R9.4.1 chemistry). Base calling, de-multiplexing, and adapter removal were performed using the Guppy (version 6.2.1+6588110a6) high accuracy base caller with a minimum quality score of 9 (default settings). The Illumina reads were trimmed and adaptors removed with fastp v0.20.1 [29] using the default settings. The ONT reads from the two rounds were merged for each isolate and processed using Filtlong v0.2.0 [30] to remove all reads shorter than 1000 bp, retaining 90% of input data (after removal of short and low-quality reads). Subsequently, the processed short and long reads were used as input into Unicycler v0.5.0 [31] in ‘normal’ mode, using default settings besides a modified start gene database that included *M. felis* and a minimum contig length of 200 bp. The assemblies where then polished using Polypolish v0.5.0 [32] using default settings, incorporating BWA-MEM v0.7.17-r1188 [33]. Full genome alignments were generated using the progressive MAUVE algorithm [34] to identify regions of reassortment and gene loss or gain.

### Genome annotation and sequencing analysis

The genomes were annotated using the Prokaryotic Genome Annotation Pipeline v2022-12-13.build6494 [35] for primary analysis. A *Tenericutes* specific protein database, containing both Swiss-Prot (4464 sequences) and TrEMBL (343,598 sequences) annotations, was downloaded from UniProt and converted to a Diamond format protein database [36]. This database was used as input to Pydeel (https://github.com/alegione/pydeel) to determine the coding ratio of our annotated genomes in comparison with *M. felis* genomes on GenBank. This tool identifies the coding ratio of annotated genes compared to their best hit within the provided database to identify the proportion of annotated genes that may have erroneous insertions or deletions as a by-product of the use of long read sequencing. From here, annotated genomes were processed through Panaroo v1.3.0 [37], along with previously sequenced *M. felis* genomes (Myco-2 and ATCC23391), to identify the core genome. The core genome results were visualised using Pagoo [38]. Gene alignments generated by Panaroo were filtered based on their presence in at least four genomes and present as a single copy per genome before analysis for genetic diversity using DnaSP v6.12.03 [39]. The annotated genomes from this study were investigated for pathogenicity islands and integrative conjugative elements using Island Viewer 4 [40] and ICE finder [41], respectively. Antimicrobial resistance genes and virulence factors were detected by generating a Diamond database of the CARD [42] and full virulence factor databases [43], and using Diamond (v 2.1.9) BLASTp to identify best hits with a cut-off bit score of 50 (all other parameters were left as their default). The UniProt webserver was used to identify homologs of specific open reading frames (ORFs) in other genomes (https://www.uniprot.org).

### Phage isolation and visualisation

A phage-like contig was identified in the annotation results of one of the assemblies (MF329). In order to confirm whether the detection of a bacteriophage genomic sequence corresponded to the presence of an integrated prophage or to replicative virions, bioinformatic methods were first utilised. The probable terminal ends were investigated using short and long read mapping to the contig with BWA-MEM and minimap2 v2.26-r1175 [44], respectively, and the contig’s starting point adjusted based on the depth of coverage. The tool PhageTerm [45] was also used to investigate the potential phage start and end points. To investigate whether this contig was a prophage, the long reads were mapped, using minimap2, to the updated phage contig. Any reads that had overhangs at the 5’ and 3’ ends of the contig were extracted and subsequently mapped to the assembled MF329 genome to determine whether there was a common point of overlap. To detect the presence of bacteriophage virions, transmission electron microscopy was performed on a culture of MF329. Isolate MF329 was cultured in 50 mL of Mycoplasma Broth, as described above, for 20 hours. The culture was centrifuged for 15 mins at 10,000 x g at 4°C and the medium fraction separated from the bacterial cell pellet. The bacterial cells were gently lysed by resuspension in 150 µL of sterile water, followed by centrifugation at 10,000 x g at 4°C for 5 mins to remove cellular debris. This was repeated twice, with the supernatant fraction captured each time for ultracentrifugation. The culture medium and the cell lysate supernatant fractions were separately ultra-centrifugated at 100,000 x g for 90 min at 4°C, and the pellets resuspended in 0.2 M HEPES buffer, pH 7.7. Negatively stained samples were examined at the Ian Holmes Imaging Centre (Melbourne, Australia) using a FEI Talos L120C cryoTEM (Talos L120C) microscope.

## Results

### Characterisation and comparisons of genomes

All four isolates were successfully assembled into circularised, complete genomes. In addition to these four genomes, a novel *M. felis* phage genome was identified that utilised the mycoplasma translation code. Metrics for long and short read sequencing output are provided in Supplementary Table 1. The assembly statistics for the four isolates are provided in Supplementary Table 2. Briefly, the circularised isolates were significantly larger than the reference genome (Myco-2; 841,695 bp), with a median length of 940,935 bp (range: 905,741 – 948,716) and a median of 759 coding domain sequences (CDSs) (range: 743 – 777), compared to the reference genome which has 740 CDSs. The NCBI Prokaryotic Genome Annotation Pipeline incorporates a CheckM [46] completeness analysis and an average nucleotide identity taxonomic classification step to identify any potential issues with an assembled genome. For all four genomes the CheckM completeness mirrored that of the reference Myco-2 genome (99.21% completeness), and all assemblies were classified as *Mycoplasma felis* with high confidence (Supplementary Table 2). The coding ratios of annotated assemblies were investigated to determine the frequency of potential indel errors. The two published *M. felis* genomes (Myco-2 and ATCC23391) had mean coding ratios of 1.0 (standard deviation ±0.02) and 0.96 (±0.32), respectively. Our completed genomes, MF47, MF219, MF329, and MF632, had mean coding ratios of 1.0 (±0.28), 0.99 (±0.30), 1.0 (±0.27), and 1.0 (±0.26), respectively. Seven recently sequenced *M. felis* isolates that utilised only Oxford Nanopore sequencing had a substantial number of indel errors in their annotations based on coding ratio analysis (mean coding ratios between 0.44 and 0.47) and were not used in further comparisons for this reason (Supplementary Figure 1).

A progressive MAUVE alignment of the four genome assemblies in comparison to that of Myco-2 (Figure 1) identified two significant genome inversions in the reference sequence compared to the isolates assembled here, as well as more localised inversions within each of the isolates. These inversions, and the genomic regions present in individual isolates, were typically flanked by transposases. Comparison of the average nucleotide identities (ANIs) of the four new assemblies, and with previously published *M. felis* genomes (Supplementary Table 3) found that the best match for each of the four Australian isolates was another isolate from this study, with the exception of MF329, noting that MF047 and MF219 were obtained from the same animal two months apart.

**Figure 1.**
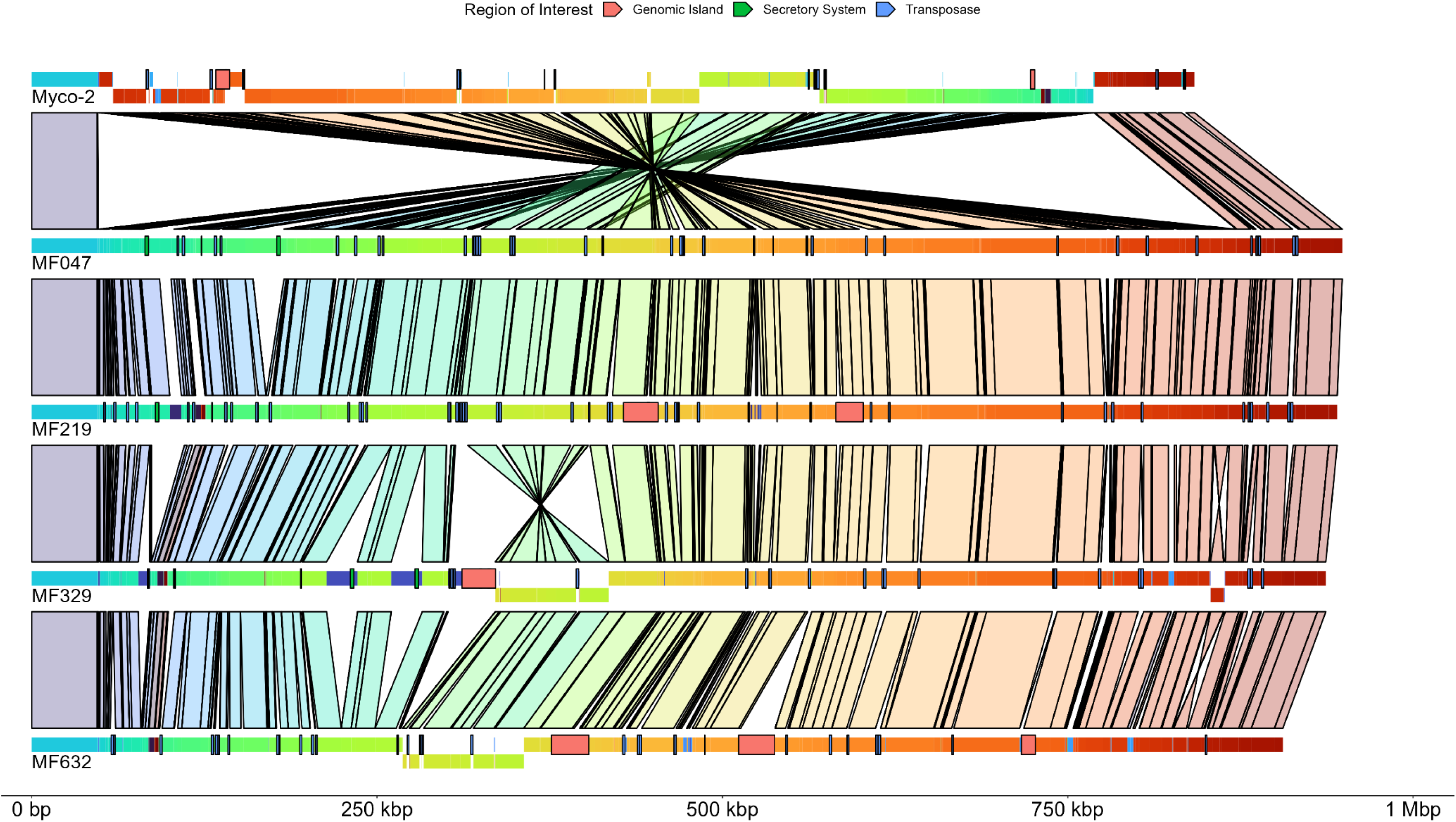
A progressive Mauve alignment of the *Mycoplasma felis* reference genome (Myco-2) and the genomes sequenced in this project. Locally co-linear blocks (LCBs) are separated by colour within genomes, and are the same colour in each genome, with links to identify placement and inversion differences. Solid blocks overlaying the genomes represent the presence of transposases, type IV secretion systems, and genomic islands, whilst solid blocks of blue that break up the colour gradient within the genomes are representative of LCBs not present in linked genomes

The core genome of *M. felis*, as determined by Panaroo (genes present in >99% of genomes) contained 594 genes, with 316 shell genes (genes present in 15% - 95% of genomes). Collapsing paralogues reduced the number of shell genes to 237. Principal component analysis (PCA) highlighted clustering of the two published genomes, as well as clustering of isolates that were obtained from the same individual (Figure 2). All genes, except those annotated as hypothetical proteins, identified in the newly sequenced isolates but not present in the reference sequence are listed in Table 2, with the homologs found in other genomes. The variations between the genomes were mostly observed in sequences encoding putative proteins involved in DNA transfer, metabolism, DNA replication, mycoplasma-host cell interactions, and restriction modification systems (Figure 3). At a gene level, an average of 2.41% of each coding sequence was a segregation site, and an average nucleotide identity per gene of 98.8% (Supplementary Table 4). Only four genes were identified as having a significant Tajima’s D value by DnaSP. These genes encoded for two hypothetical proteins (Myco-2 loci 000416 and 000586), 23S rRNA (pseudouridine^1915^-*N*^3^)-methyltransferase (Myco-2 locus 000375), and a Sua5/YciO/YrdC/YwlC family protein (Myco-2 locus 000629). Each of these coding regions had a Tajima’s D value below 1, suggesting neutral selection (Supplementary Table 4).

**Figure 2.**
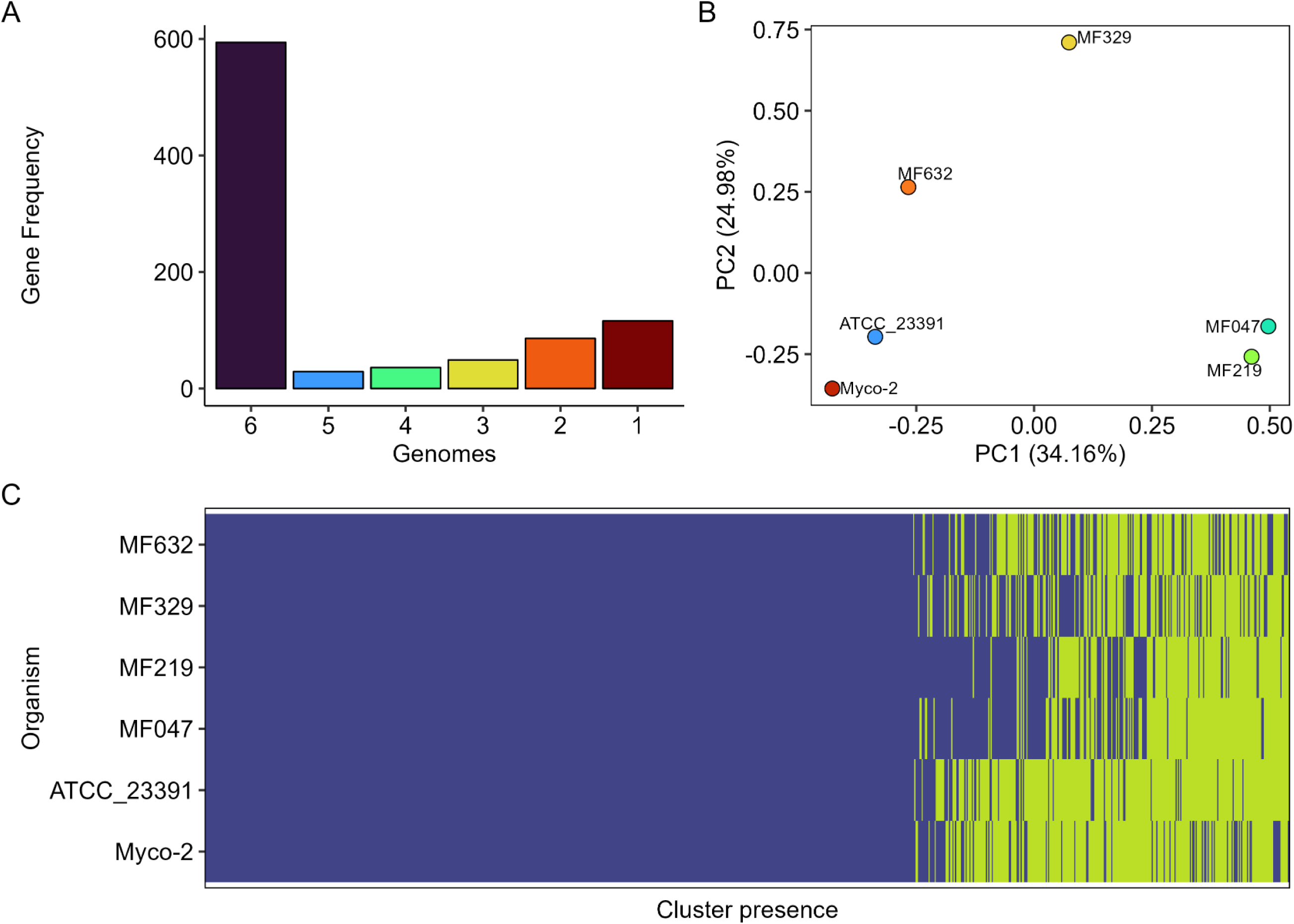
Analysis of the *Mycoplasma felis* core genome and gene presence/absence using Panaroo and plotted using Pagoo. A) the frequency of clustered genes within the six genomes used in the analysis; B) principal component analysis of gene presence/absence using Bray Curtis dissimilarity identifies clustering of the previously published genome sequences; C) clusters of genes present (blue) or absent (yellow) in the six genomes included in analysis.

**Figure 3.**
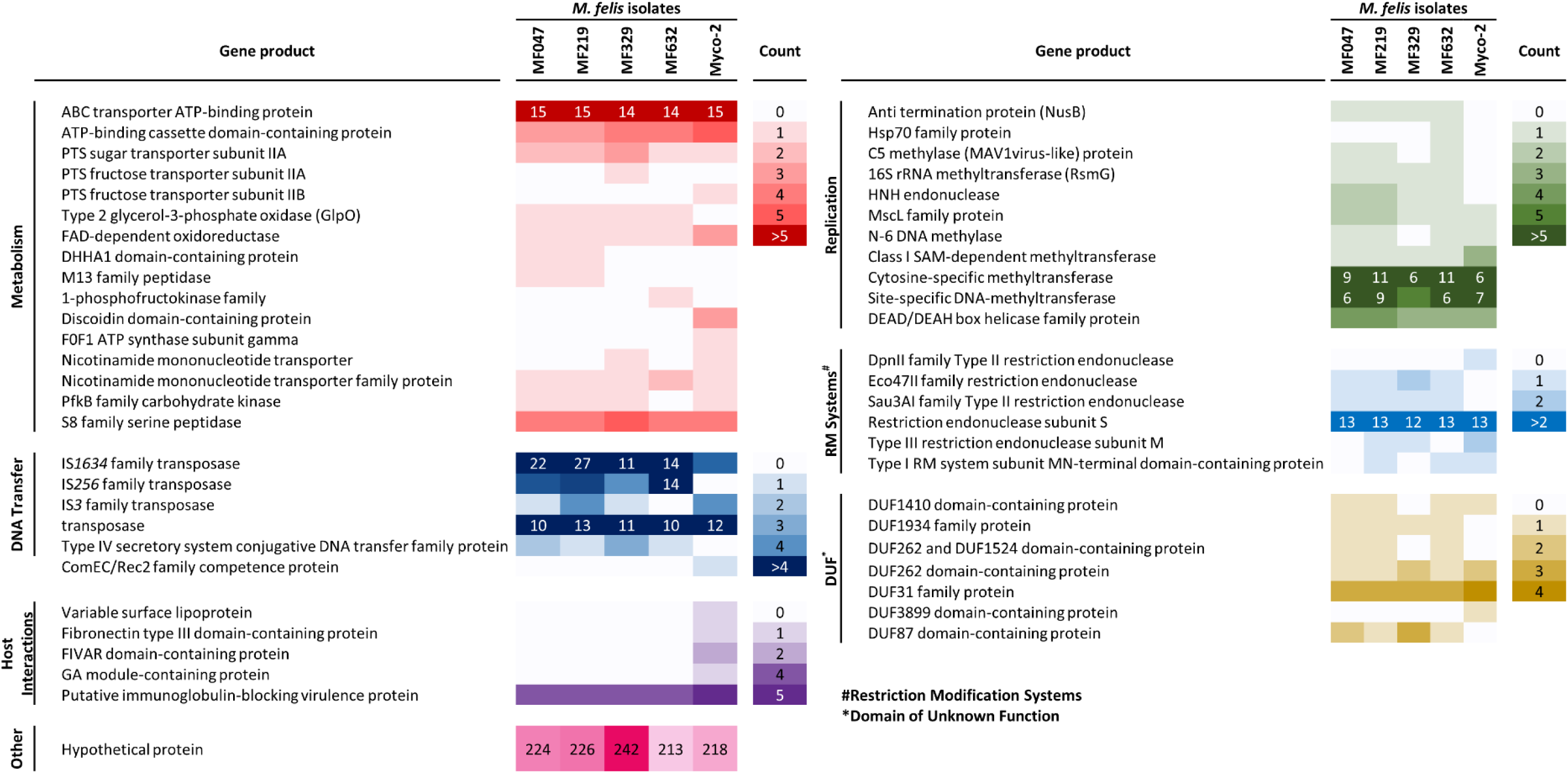
Presence and absence of genes within the genomes of Australian felid *M. felis* isolates compared to the genome of the Myco-2 equid *M. felis* reference strain, clustered by putative protein functions. Only genes that differ between isolates or differ from the reference are listed. Counts indicate the number of gene copies detected within each isolate.

**Table 2.** Coding sequences detected in the genomes of Australian *M. felis* isolates but absent in the equid *M. felis* reference strain (Myco2), and their possible species of origin based on amino acid sequence identity.

| Coding sequence | Possible species of origin | % Amino acid pairwise<br>sequence identity | Predominant host |
| --- | --- | --- | --- |
| Type IV secretory system conjugative DNA transfer family<br>protein | *M. cynos* | 43.2 | Dog |
| IS256 family transposase | *M. penetrans* | 50.9 | Human |
| Type 2 glycerol-3-phosphate oxidase (GlpO) | *M. canis* | 87.2 | Dog |
| PTS fructose transporter subunit IIA | *M. flocculare* | 52.6 | Swine |
| DHHA1 domain-containing protein | *M. canis* | 84.8 | Dog |
| M13 family peptidase | *M. canis* | 65.4 | Dog |
| 1-phosphofructokinase family | *M. canis* | 52.6 | Dog |
| Anti-termination protein NusB | *M. mustelae* | 61.6 | Mink |
| Hsp70 family protein | *M. flocculare* | 85.4 | Pig |
| C5 methylase (MAV1virus-like) protein | *M. columboralis* | 70.2 | Pigeon |
| 16S rRNA methyltransferase (RsmG) | *M. mustelae* | 63.3 | Mink |
| HNH endonuclease | *M. columbina* | 38.0 | Pigeon |
| Eco47II family restriction endonuclease | *M. mycoides* subsp. *mycoides* SC | 49.6 | Cattle |
| Sau3AI family type II restriction endonuclease | *M. crocodyli* | 52.7 | Crocodile |

The genomes of the newly sequenced isolates had 1-3 secretory systems (Type IV secretory systems), whilst that of Myco-2 has no annotated secretory systems. The newly annotated genomes also had an increased number of transposases, typically annotated as IS*1634* or IS*256*, or as IS*3*. In comparison, the Myco-2 genome only had IS*3* and IS*1634* family transposase annotations.

### Genomic islands, virulence genes and antimicrobial resistance genes

Genomic islands were detected in three of the four newly sequenced genomes (**Error! Reference source not found.**). The average length of these islands was 22,145 bp (± 5818 bp) and each contained an average of 16 (±5) annotated genes. In comparison, the Myco-2 strain of *M. felis* had two predicted genomic islands of 3052 bp and 10,027 bp. Further inspection of these predicted genomic islands identified matching regions present in isolates in alternative locations. For example, MF329 had a predicted genomic island of 24 kb (nucleotide position [nt] 311,409 – 335,810) containing 19 genes, including a type IV secretion system and an insertion sequence (IS*1634*) at its midpoint. Extraction and remapping of the sequence of this genomic island showed that it was repeated three times within a wider 122 kb region (nt 213,995 – 335,810) of the MF329 genome. This same genomic region occurred, without the IS element present, in the genomes of MF047 (twice), MF219 (once), and MF632 (once). In the case of MF632, the region was also a predicted to be a genomic island, but in this instance was flanked with restriction endonuclease genes and either predicted transposases (3’) or an IS*256* element (5’).

Diamond BLAST searches for antimicrobial resistance genes (CARD database) and virulence genes (Virulence Factor Database) identified few high identity hits (Table 4 and Supplementary Tables 5 and 6). Only one hit with the virulence factor database had an amino acid sequence identity of greater than 90%, and this was to the gene for the elongation factor Tu in *Mycoplasma synoviae* (90.1% average identity), which has been found to be an adherence factor and is present in all genomes in this study. Only five genes had hits with >50% amino acid sequence identity. The closest hits to these genes were *oppF* (59.9% identity; *Mycoplasma mycoides* subsp. *mycoides*; immune modulation), *eno* (58.9%; *Streptococcus pneumoniae*; exoenzyme), *gap* (57.9%; *Mycoplasma genitalium*; adherence), *dnaK* (53.9%; *Chlamydia trachomatis*; adherence), and *sugC* (50.4%; *Mycobacterium marinum*; nutritional/metabolic factor). No genes had hits with >50% amino acid sequence identity with antimicrobial resistance genes in the CARD database.

**Table 3.** The location of predicted genomic islands in the genomes of *M. felis* isolates characterised in this study, as well as the reference genome Myco-2.

| Strain | Genomic island | Start – End<br>(relative to genome) | Length | Total number of genes<br>(non-hypothetical) |
| --- | --- | --- | --- | --- |
| Myco-2 | 1<sup>b,d,e</sup> | 133250 - 143277 | 10027 | 11 (5) |
|  | 2<sup>b,c,d,e</sup> | 723039 - 726091 | 3052 | 8 (6) |
| MF047 | - | - | - | - |
| MF219 | 1<sup>a,b,d,*</sup> | 428340 - 453515 | 25175 | 15 (7) |
|  | 2<sup>a,b,d,e</sup> | 582071 - 602042 | 19971 | 11 (6) |
| MF329 | 1<sup>b,c,#</sup> | 311409 - 335810 | 24401 | 19 (15) |
| MF632 | 1<sup>a,b,d,*</sup> | 376254 - 403316 | 27062 | 19 (8) |
|  | 2<sup>b,#</sup> | 511813 - 537933 | 26120 | 22 (15) |
|  | 3<sup>a,b,c,d</sup> | 716455 - 726594 | 10139 | 8 (1) |

<sup>a, b, c, d, e</sup> – Predicted genomic island is present with some variation, such as the presence/absence of a transposase, in (a) Myco-2, (b) MF047, (c) MF219, (d) MF329, or (e) MF632, but not predicted to be a genomic island in those cases.
<sup>\*, #</sup> - Predicted genomic islands with structural similarity across different genomes
Genomic islands determined with the Island Finder 4 web tool. None were detected in the MF047 isolate.

**Table 4.** Quantity and range of amino acid sequence identities and similarities of putative virulence and antimicrobial resistance genes (AMR) based on significant BLASTP hits (bit scores >50) compared to the virulence factor database and CARD database, respectively.

| Genome | Putative virulence genes |  |  | Putative AMR genes |  |  |
| --- | --- | --- | --- | --- | --- | --- |
|  | Total | Median identity (min – max) | Median similarity (min – max) | Total | Median identity (min – max) | Median similarity (min – max) |
| Myco-2 | 48 | 31.7 (24.4 - 90.1) | 52.6 (44.4 - 95.9) | 24 | 29.8 (22.5 - 40.1) | 51.7 (39.1 - 69.6) |
| MF047 | 49 | 31.6 (24.6 - 90.1) | 52.7 (41.8 - 95.9) | 25 | 29.4 (22.6 - 40.3) | 50.6 (39.2 - 69.6) |
| MF219 | 49 | 31.6 (24.6 - 90.1) | 52.4 (41.8 - 95.9) | 25 | 29.6 (22.6 - 40.3) | 50.6 (39.2 - 69.6) |
| MF329 | 50 | 32 (24.4 - 90.1) | 52.2 (41.5 - 95.9) | 25 | 29.7 (22.5 - 40.2) | 50.6 (39.1 - 69.6) |
| MF632 | 51 | 31.6 (24.4 - 90.1) | 52.4 (41.5 - 95.9) | 26 | 29.7 (22.4 - 40.2) | 51 (38.9 - 69.6) |

### Phage detection and identification

A 33.6 kb contig (vB Mfe PM329; *<u>M</u>ycoplasma <u>fe</u>lis* strain <u>PM329</u>) was assembled and identified as a likely phage or prophage in the sequenced *M. felis* MF329 isolate. It had an average depth of coverage of 40,847.7 reads, with 82.6% of the MF329 Illumina reads mapping to the phage genome. Interrogation of the Nanopore read dataset to determine its genome integration status detected only 63/18134 (0.35%) reads that mapped to both the phage and the bacterial genome. Of these, most were found to map across the bacterial genome at mostly single-read depth, with the exception of five reads that clustered mid-genome in the region of nt 456,500. If this was indicative of a true prophage insertion site it would disrupt ORF MF329_000403, which encodes a putative 16S rRNA pseudouridine synthase gene. The phage genome had a 27.1% G+C content and encoded 36 predicted ORFs using the mycoplasma translation code. It shared 94.8% global pairwise nucleotide sequence identity with two contigs in the publicly available unassembled *M. felis* contig dataset (NCBI Biosample: SAMN30346842), and there was 96.8% and 97.4% pairwise amino acid sequence identity between each of their encoded putative recombinase and portal proteins, respectively. Interrogation of the read datasets of the other three isolates sequenced in this study with the phage genome identified only 4/9515 (0.04%; MF047), 19/19128 (0.1%; MF632) and 22/65383 (0.03%; MF219) mapped Nanopore reads, within the range of barcode error. No Illumina reads mapped to the phage genome, indicating that the isolates did not carry the phage. Phylogenetic analyses using VipTree [47] and vContact2 [48] was not able to assign vB Mfe PM329 to any known phage genus or family, but its gene arrangement was similar to those of the MAgV1 phage from *M. agalactiae* and MAgV1-like prophage sequences detected in mycoplasma species in the Hominis phylogenetic group. In particular, both the protein and gene arrangement similarity were greatest to those of prophages found in *Mycoplasma mustelae* and *Mycoplasma molare*, including the central position of the recombinase gene. BLASTX analysis of predicted ORFs detected pairwise protein sequence identities of 48% to 69% between core conserved ORFs encoded within the *M. mustelae* and *M. molare* prophage sequences [49] (Supplementary Table 7).

Confirmation of productive virus formation and, therefore, active replication, was achieved using cryoTEM on medium and cell lysate supernatant fractions, which revealed the presence of non-enveloped contractile-tailed polyhedral viral particles approximately 100 nm in length, with a 50 nm diameter head and tails of varying lengths. Higher numbers of virions were observed within the cell lysate supernatant fraction than in the culture medium fraction, with clusters of virions observed seemingly embedded within bacterial membrane debris (Figure 4). These long tailed polyhedral viral structures were consistent with those seen in siphoviruses and myoviruses within the recently ratified *Caudoviricetes* class of bacteriophage. The contractile tail and polyhedral head structure is consistent across all visualised mycoplasma phages, with the main differences being the head diameter and tail lengths [50][51].

**Figure 4.**
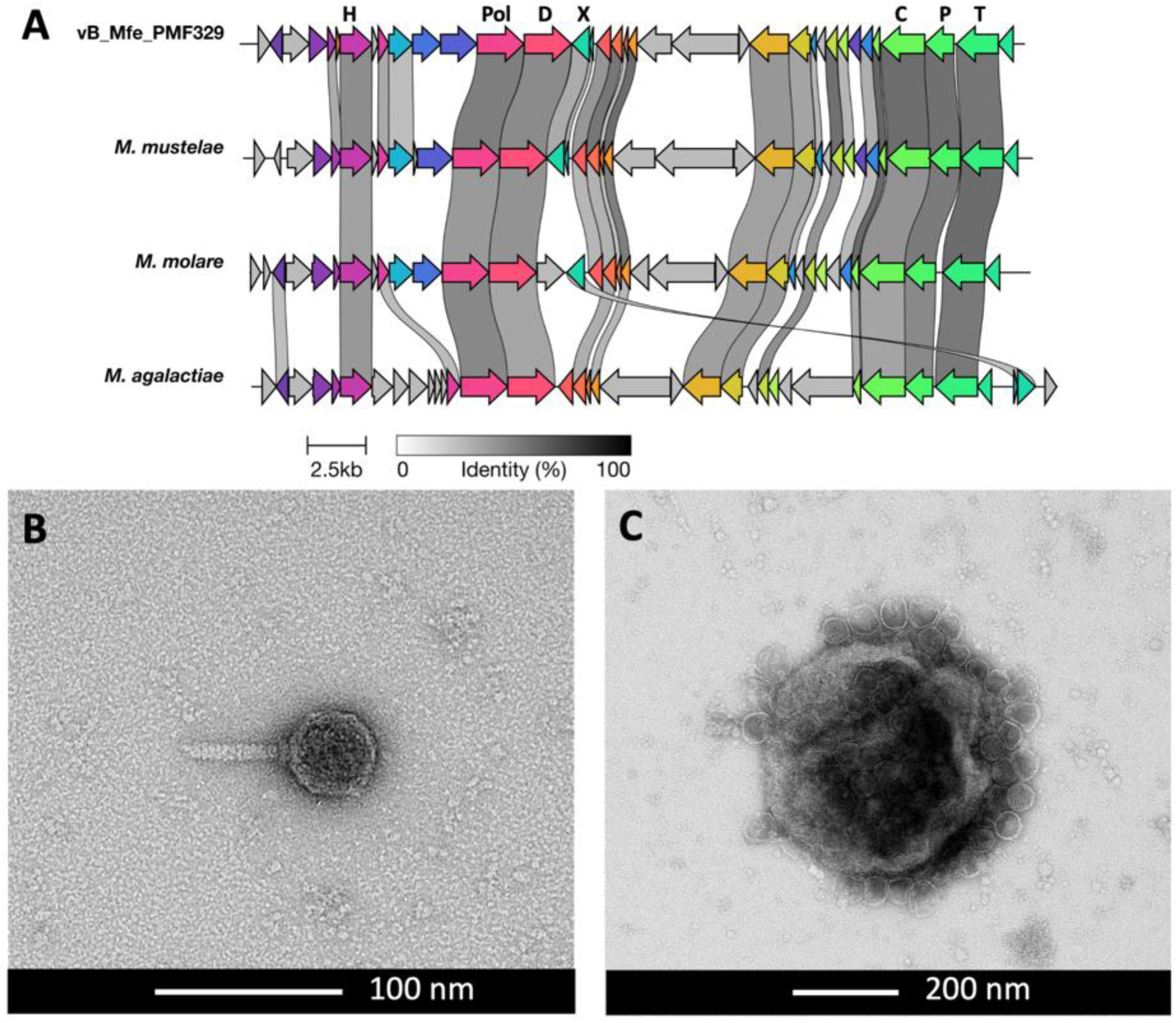
A) Genome of the bacteriophage identified in *Mycoplasma felis* strain MF329 compared to MAgV1-like prophage sequences found in *M. mustelae*, *M. molare* and *M. agalactiae*. Genome annotations depicting conserved phage/prophage gene sequences are indicated in black. H = helicase; Pol = DNA polymerase; D = DNA primase; X = Xer recombinase; C = prohead protein; P = portal protein; T = terminase; HP = hypothetical protein. B) and C) cryo-transmission electron photomicrographs of bacteriophages detected in a cell lysate of *M. felis* isolate MF329, showing that the phage structure consists of a contractile tail and polyhedral head.

## Discussion

In this study, the genomes of four new *M. felis* isolates were characterised and compared; three of the isolates were derived from clinical cases of respiratory disease and the remaining one from an unresolved post-surgical infection. Comparisons between these genomes and with several published datasets derived from other isolates of *M. felis* detected genomic variations and inversions between the Australian felid isolates and the only assembled *M. felis* reference isolate, from a horse. These variations and inversions were flanked by transposases, which are commonly involved in HGT and genome reassortment in other mycoplasmas. There were differences in the number of copies of putative transposases from several families, including the IS*1634*, IS*256* and IS*3* (Figure 3). Insertion sequences (ISs) are transposable elements that are often present in multiple copies in a genome and involved in chromosomal rearrangements by deletion, insertion, or inversion. The substantial similarity between various isoforms of ISs enables them to facilitate homologous recombination [52]. IS families encoded in mycoplasma genomes play an important role in genomic plasticity with several reported instances of IS exchange between strains or species [15, 53–56]. This is in contrast to early research that suggested that some ISs, such as IS*1634*, which was found in abundance in the sequenced genomes of our isolates, were unique to specific mycoplasma species [57]. The differences between our Australian *M. felis* genomes, which were obtained from three different cats, based on gene presence/absence and the number of coding sequences, underscores the genetic diversity present amongst the sequenced felid *M. felis* isolates (Figure 2). Multiple gene insertions, relative to the equid *M. felis* reference genome, were detected within the cluster of Australian field isolates (Table 2). These included genes involved in DNA transfer, replication, metabolism, and restriction modification (RM) systems, as well as a number of proteins of unknown function (Figure 3).

### Gene gains compared to the equid M. felis reference strain

Of particular interest are the gene acquisitions involved in DNA transfer, such as the multiple copies of genes encoding putative type IV secretory system (T4SS) conjugative DNA transfer family proteins within the Australian felid genomes, which were not detected in the publicly available equid and felid *M. felis* genomes. T4SS conjugation systems translocate DNA through a process dependent on cell to cell contact and have been identified as one of the core genes within mycoplasma Integrative and Conjugative Elements (ICEs), which play a significant role in genomic plasticity in the species in which they have been found [58, 59]. ICEs have been shown to be self-transmissible in *M. agalactiae*, providing conjugative properties from ICE-positive cells to ICE-negative cells, while concurrent chromosomal transfers (CTs) occur in the opposite direction [60]. Protein sequence analysis of the putative T4SS identified in the Australian felid *M. felis* isolates detected 43.2% amino acid sequence identity with ORF5 of the ICE in *Mycoplasma cynos*. Whilst *M. cynos* predominantly infects the respiratory tract of dogs, it has also been detected in cats in association with conjunctivitis and upper respiratory tract disease [61]. Taken together, it is plausible that a potential HGT event may have resulted in transfer of the T4SS conjugative element of an ICE from a *M. cynos*-like species to *M. felis* in a shared feline host. This T4SS was part of a defined region of genes that, along with 15 hypothetical proteins and a DUF87 domain-containing protein, was repeated three times in the genome of isolate MF329. DUF87 domain-containing proteins are associated with conjugative elements in other organisms, and include TraJ in the virulence plasmid of *Salmonella enterica* serovar Typhimurium [62], adding further weight to the likelihood that this region is an ICE, even though the bioinformatics tools used here did not detect it.

Several genes involved in metabolism were newly detected in the felid *M. felis* isolates sequenced in this study. A type 2 glycerol-3-phosphate oxidase (GlpO) was identified within the Australian *M. felis* genomes, while in the equid *M. felis* isolate, only its FAD-dependent oxidoreductase domain was detected, accompanied by two transposases, located upstream. GlpO converts glycerol-3-phosphate to dihydroxyacetone phosphate (DHAP), releasing hydrogen peroxide as a by-product, which has been shown to play a role in cytotoxicity *in vitro* in *M. gallisepticum* and *Mycoplasma pneumoniae* [63, 64], and in the virulence of *M. mycoides* subsp. *mycoides* small colony type and mosquito-associated *Spiroplasma* species [65, 66]. The significant reduction or complete loss of H2O2 production in highly passaged *M. bovis*, *M. agalactiae* and *M. gallisepticum* strains, and in attenuated vaccine strains of *M. gallisepticum*, compared to wild type strains suggests that glycerol metabolism can rapidly become redundant in some mycoplasmas [63, 67, 68]. It is possible that the sequences coding for GlpO were lost in the equid *M. felis* isolate, possibly because of differing levels of available glycerol in horses compared to cats. Likewise, variations were seen in the counts of genes coding for nutrient transport systems, including ATP-binding cassette (ABC) transporters and the PEP-dependent phosphotransferase transport system (PTS), as well as numerous other metabolic enzymes. A number of these nutrient transport systems, and many cell surface enzymes, are multifunctional in mycoplasmas, playing critical roles in pathogenesis and survival in the host [13, 69–74]. The differences observed in the PTS and ABC transporters of the Australian felid *M. felis* isolates, in contrast to the equid *M. felis* isolate, may also be indicative of rapid metabolic gene gains and losses in response to the varying nutrient availabilities within felid and equid ecological niches.

In the Australian felid *M. felis* genomes, five new coding sequences related to replication were identified that were absent in the equid origin *M. felis* genome, including genes encoding the putative anti-termination protein NusB, a Hsp70 family protein, a C5 methylase, the 16S rRNA methyltransferase RsmG, and an HNH endonuclease. The presence of these additional proteins suggests potential differences in bacterial growth within the feline and equine hosts, potentially due to distinct replication strategies. Future experiments comparing growth patterns of these isolates in felid or equid derived cells are indicated. The potential origins of these additional sequences have been detected in mycoplasmas isolated from mink, pigs, and dogs, which share a common ecological environment with cats, or pigeons, which are a common prey of cats. The HGT events resulting in the acquisition of these genes by the Australian felid *M. felis* isolates may have been facilitated by the close ecological proximity or may have occurred within the feline gastrointestinal tract (GI) following the ingestion of infected prey. HGT is frequently observed between diverse microorganisms within the mammalian gastrointestinal tract, possibly due to the high population density in this niche, which presumably facilitates genetic exchange [75]. Furthermore, the conjugation mechanism favoured for HGT within the GI tract protects DNA from nucleases and heavy metals, and thus is more likely to result in a successful genetic exchange [76].

Two additional restriction endonucleases (Eco47II and Sau3AI) were annotated in the Australian felid *M. felis* genomes, and they lost a DpnII family type II restriction endonuclease, compared to the equid isolate. Restriction modification (RM) systems play a pivotal role in regulating bacterial HGT by protecting the genome against incoming mobile genetic elements, or by facilitating recombination with the introduced DNA [77]. A strong correlation has been reported between the presence of active Type II RM systems and the presence of mobile genetic elements and ICEs within *M. agalactiae* genomes, indicating the facilitative role of Type II RM systems in HGT in mycoplasmas [78]. The diversification of RM target recognition sites can regulate HGT by promoting genetic exchanges among those organisms with RM systems that recognise the same target motifs, and by reducing genetic transfer between lineages that possess distinct RM systems [79]. Taken together, it is plausible that following a host switch into horses, *M. felis* may have diversified its RM systems to enhance the level of genetic exchange from phylogenetically distant bacteria.

### Gene losses compared to the equid M. felis reference strain

A sequence encoding a putative ComEC/Rec2 family competence protein was absent in the Australian felid *M. felis* isolates. This protein is one of the components of DNA translocation via transformation, enabling DNA transport by forming a channel across the cytoplasmic membrane [80]. While the ComEC component has been reported to be essential for DNA transport in Gram positive bacteria, deletion of homologous genes did not affect DNA uptake in bacteria without a cell wall [81]. Furthermore, the competence domain seems to be missing in many mycoplasmas, possibly due to the reductive evolution of these bacteria [82]. Characterisation of two mycoplasma genomes recovered from gut microbiota of a deep-sea isopod detected the *comEC* gene adjacent to genes encoding enzymes producing dTMP from thymidine or dUMP, suggesting that the role of the ComEC protein might be importing extracellular DNA as a nutrient source [83]. The presence of ComEC in the equid *M. felis* isolate, and its absence in the Australian felid *M. felis* isolates suggests that during evolutionary adaptation DNA translocation via a membrane channel might have become redundant and perhaps was replaced with T4SS conjugation-mediated DNA transfer, as discussed above.

The Australian felid *M. felis* genomes exhibited a total absence of four coding sequences associated with host cell interactions, those encoding a putative variable surface lipoprotein, a fibronectin type III domain-containing protein, a FIVAR domain-containing protein, and a GA module-containing protein, in contrast to the equid *M. felis* genome. The FIVAR (Found In Various Architectures Region) and GA domains are essential constituents of extracellular matrix-binding proteins (Embp), facilitating biofilm formation and attachment to fibronectin in some bacteria such as *Staphylococcus epidermidis* and *S. aureus* [84, 85]. The presence of more than 30% amino acid sequence identity between the FIVAR and GA domains in the equid *M. felis* reference isolate Myco-2 and these domains in Embp of *S. aureus* raises the possibility that the equid *M. felis* isolate might have potential for biofilm formation and/or fibronectin attachment. Conversely, the absence of these proteins, and of the surface lipoprotein and fibronectin type III domain-containing protein, suggests that these capacities may have become obsolete in the Australian felid *M. felis* isolates, possibly indicating different modes of host cell interaction in the two hosts.

### Rapid evolution within two months in one host

Two of the Australian *M. felis* isolates sequenced in this study, MF047 and MF219, were obtained from separate bronchoalveolar lavages (BAL) of a feline patient with clinical signs consistent with pneumonia. Isolate MF047 was obtained from a BAL collected during the initial consultation, whereafter the cat was treated with 25 mg of doxycycline twice a day orally for approximately two months. Because the response to treatment was unsatisfactory, a second BAL was collected and isolate MF219 was obtained. While no genome reassortment was detected between the MF047 and MF219 genomes, minor differences were observed in the counts of some genes encoding proteins involved in RM systems, DNA transfer, and methylation (Figure 3). These variations may suggest rapid host adaptation of *M. felis* after only two months. The growing utilisation of whole genome sequencing in mycoplasma diagnostics is revealing links between genetic mutations and antimicrobial resistance. Genomic comparisons of MF047 and MF219 revealed the presence of five single nucleotide polymorphisms (SNPs) within the gene encoding the ribosomal RNA small subunit methyltransferase A (RsmA/KsgA), along with one SNP in the gene encoding the 16S rRNA and another SNP in the gene encoding the 23S rRNA. In other studies, mutations within the 16S RNA methyltransferase family have been reported to confer aminoglycoside resistance in *M. bovis* and *M. gallisepticum* [86, 87], while mutations in the 16S rRNA or 23S rRNA genes have been shown to be associated with resistance to tetracyclines and macrolides, respectively, in *M. pneumoniae*, *M. hominis*, *M. genitalium*, and *M. bovis* [86, 88–90]. MF047 and MF219 both contained five protein coding regions with low confidence matches to tetracycline resistance genes, with the highest bit score of the five associated with a predicted elongation factor G protein with amino acid sequence identity/similarity to *tet*(T) from *Streptococcus pyogenes* of 25.8%/47.7%. It is important to emphasise that the application of novel genome sequencing techniques for mycoplasma diagnostics, particularly in the context of antimicrobial resistance, is still in its early stages, and the availability of data for comprehensive analysis of all mycoplasma species (and particularly animal mycoplasmas) is currently limited. Future research determining the phenotypic susceptibility of isolates MF047 and MF219 to tetracyclines is necessary to determine whether there is a correlation between the genomic variations observed and tetracycline resistance.

### Phage in mycoplasmas

In other bacteria, phages have been demonstrated to provide critical virulence factors to their hosts, and play a key role in the pathogenesis of diseases associated with a diverse range of bacteria (e.g. cholera, diphtheria, botulism, and haemolytic-uremic syndrome). Approximately 20 mycoplasma phages/prophages have been reported across diverse ruminant, avian, canine, pig, human and rodent mycoplasmas [49, 91]. Of these, most are unclassified, only three have had their virion structure visualised (P1, Hr1 and Br1), and only one has been classified by the International Committee for the Taxonomy of Viruses (ICTV), the temperate *Mycoplasma pulmonis* phage P1 (12 kb genome), as the sole member of the floating genus *Delislevirus* [92]. The well-described temperate MAV1 phage found in *Mycoplasma arthritidis* (16 kb genome) has not yet (at the time of publication) been classified within the new system. In other mycoplasmas, phages have demonstrated dynamic mobility within the bacterial chromosome, with loss of integrated prophages seen with clonal passage (MFV1 in *Mycoplasma fermentans*), and variable expression of unique phage-encoded membrane surface proteins (MFV1 and MAV1 in *M. fermentans* and *M. arthritidis*, respectively) [93], and may have a role in bacterial adaptation and survival. In *M. felis*, vb MFe PM329 appears to also be a temperate phage, capable of being virion-productive and potentially have a lysogenic phase. No known selective advantage or disadvantage of the presence of this phage within *M. felis* was able to be determined in this study because of the low number of *M. felis* genomes available. Detection of a contig similar to that of vb MFe PM329 within *M. felis* 16-057065-4 (NCBI Biosample: SAMN30346842) from Canada indicates that this phage is probably carried by *M. felis* strains globally, although in that study it was not identified as a phage nor prophage. Future epidemiological studies on isolates from healthy and clinical disease cases are indicated to investigate the potential role of phage carriage in feline mycoplasma-associated disease.

## Conclusions

This study has provided a comprehensive analysis of the first high-quality, complete genomes of *M. felis* isolated from cats, and characterised a novel mycoplasma phage. It provides valuable insights into the intricate genomic dynamics of *M. felis*, showcasing a complex interplay of gene gains and losses, horizontal gene transfer events, and adaptations to new host environments. These findings suggest divergent evolutionary paths in different hosts and offer compelling avenues for further research into the evolutionary mechanisms and ecological factors that influence the genomic diversity of mycoplasmas. Such studies include investigating the variation in metabolism, replication, virulence, antimicrobial resistance and other functional properties in the strains with and without different genes of interest highlighted here. Additionally, the significance of gene gains/losses, and the presence of the newly identified phage, in relation to niche adaptation and host specificity requires targeted studies focusing on collecting sufficient *M. felis* samples from clinical equine and feline cases.

## Author contributions

R.N.B. performed the bacterial isolation and initial classification; A.R.L., S.M.K. and P.K.V. prepared the DNA samples, performed the sequencing and analysis; P.K.V. isolated the phage for TEM; P.K.V., S.M.K., G.F.B. and A.R.L. conceptualised the study design; P.K.V., S.M.K. and A.R.L. acquired the funding; S.M.K., A.R.L. and P.K.V. drafted the manuscript, and R.N.B. and G.F.B. revised the manuscript.

## Acknowledgments

The authors would like to gratefully acknowledge the small animal medicine team of the U-Vet Werribee Animal Hospital at The University of Melbourne for providing the samples taken from clinical cases, and Eric Hanssen at the Ian Holmes Imaging Centre for the cryoTEM imaging of the vB Mfe PM329 phage.

