## Supplementary Tables 1-3, 7, Supplementary Figure 1 for "Unveiling genome plasticity and a novel phage in *Mycoplasma felis*: genomic investigations of four feline isolates"

### Supplementary Data

**Supplementary Table 1.** Summary of DNA sequencing results after trimming and quality filtering

| | MF047 | MF219 | MF329 | MF632 |
|---|---|---|---|---|
| **Illumina reads** | | | | |
| Total paired reads | 8,977,712 | 6,647,214 | 6,481,506 | 6,807,888 |
| Median length | 151 | 151 | 151 | 151 |
| >Q20 (%) | 96.93 | 95.7 | 96.83 | 96.99 |
| >Q30 (%) | 91.31 | 88.43 | 91.12 | 91.41 |
| GC (%) | 24.64 | 24.67 | 26.35 | 24.85 |
| **Nanopore reads**[‡] | | | | |
| Total reads | 65,383 | 9,515 | 103,977 | 19,128 |
| Median length | 7,049 | 6,563 | 3,717 | 5,058.5 |
| N50 | 16,885 | 22,105 | 12,331 | 16,399 |
| >Q20 (%) | 48.38 | 51.69 | 41.64 | 49.27 |
| >Q30 (%) | 15.12 | 16.77 | 11.64 | 15.74 |
| GC (%) | 24.9 | 24.79 | 27.19 | 25.08 |

[‡]Results presented are from combining two nanopore sequencing runs

**Supplementary Table 2.** Summary of assemblies and annotations of four isolates from this study, compared to the complete reference sequence for *Mycoplasma felis* (Myco-2)

| Isolate | Myco-2 | MF047 | MF219 | MF329 | MF632 |
|---|---|---|---|---|---|
| **Accession** | NZAP022325 | CP114890 | CP114889 | CP115656 | CP114888 |
| **Assembly information** | | | | | |
| Length (bp) | 841,695 | 948,716 | 945,056 | 936,813 | 905,741 |
| CheckM completeness (%) | 99.21 | 99.21 | 99.21 | 99.21 | 99.21 |
| PGAP ANI* | 98.4% | 98.5% | 98.6% | 98.6% | 98.5% |
| **Median sequencing depth** | | | | | |
| Short reads | - | 1762 | 1372 | 467 | 1499 |
| Long reads | - | 695 | 110 | 88 | 181 |
| **Genomic features** | | | | | |
| CDS | 740 | 759 | 777 | 759 | 743 |
| ncRNA | 2 | 2 | 2 | 2 | 2 |
| rRNA | 9 | 9 | 9 | 9 | 9 |
| Regulatory | 2 | 2 | 2 | 2 | 2 |
| tRNA | 29 | 30 | 29 | 29 | 30 |
| tmRNA | 1 | 1 | 1 | 1 | 1 |
| **Genes of interest** | | | | | |
| Secretory Systems | 0 | 2 | 1 | 3 | 1 |
| Transposase | 22 | 40 | 52 | 27 | 38 |

* average nucleotide identity to *Mycoplasma felis* ATCC 23991 according to the internal PGAP ANI taxonomic classification

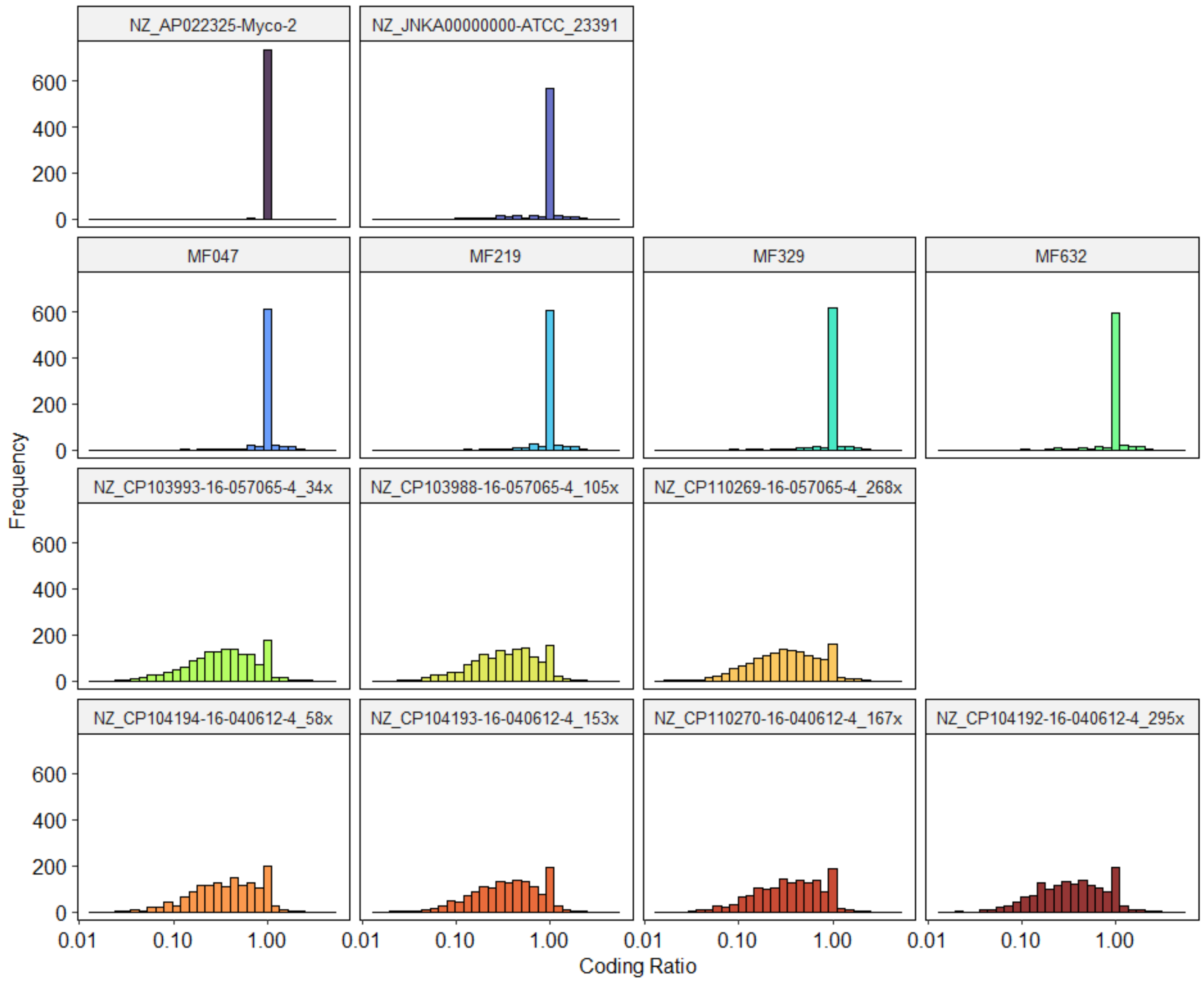


**Supplementary Figure 1.** Coding ratio frequency counts across the *Mycoplasma felis* genomes from this study and those currently available on GenBank. Coding ratios were determined by dividing the amino acid sequence length of annotated proteins by the best hit match in a custom UniProt Mycoplasma database. The coding ratio axis is log transformed for visualisation. Title names for NCBI origin data include the accession number and strain name (e.g. NZ_CP104194, and 16-040612-4). For Oxford Nanopore data from NCBI, the depth of coverage is added from NCBI (e.g. 58 x).

**Supplementary Table 3.** Average nucleotide identities (ANI) of Australian *M. felis* isolate genomes, the genome of the equid *M. felis* reference strain (Myco-2) and felid *M. felis* contig datasets (Genbank accessions: MF047: CP114890; MF219: CP114889; MF329: CP115656; MF632: CP114888; Myco-2: AP022325; 16-057065-4: CP103988; 16-040612-4:CP104192; ATCC23391: JNKA01000000).

| | MF047 | MF219 | MF329 | MF632 | Myco-2 | 16-057065-4 | 16-040612-4 |
|---|---|---|---|---|---|---|---|
| **MF219** | 99.79 | - | - | - | - | - | - |
| **MF329** | 98.21 | 98.16 | - | - | - | - | - |
| **MF632** | 98.21 | 98.17 | 98.33 | - | - | - | - |
| **Myco-2** | 98.14 | 98.06 | 98.18 | 98.17 | - | - | - |
| **16-057065-4** | 97.77 | 97.80 | 98.25 | 97.77 | 97.77 | - | - |
| **16-040612-4** | 97.65 | 97.65 | 97.84 | 97.80 | 97.72 | 97.65 | - |
| **ATCC23391** | 98.31 | 98.29 | 98.39 | 98.27 | 98.33 | 97.95 | 97.95 |

**Supplementary Table 7. Phage ORF table excluding hypothetical proteins. BLASTx amino acid pairwise identity is shown for conserved coding sequences found within other mycoplasma species.**

| Min | Name | Length | *M. mustelae* | *M. molare* | *M. agalactiae* | *M. bovis* |
|---|---|---|---|---|---|---|
| 1293 | HNH endonuclease | 528 | 41% | 57% | 56% | 53% |
| 2987 | Metallo-hydrolase | 807 | 38% | 40% | 32% | 33% |
| 3775 | Endonuclease | 357 | 51% | 51% | 47% | 47% |
| 4303 | DNA Helicase | 1368 | 53% | 49% | 48% | 48% |
| 7406 | Methionine adenosyltransferase | 1209 | - | 42% | - | - |
| 8607 | DNA cytosine methyltransferase | 1554 | 34% | 34% | 35% | - |
| 10172 | DNA polymerase | 2010 | 59% | 52% | 49% | 49% |
| 12194 | DNA Primase | 2016 | 54% | 48% | 42% | 42% |
| 14231 | Xer Recombinase | 750 | 44% | 36% | 35% | 36% |
| 24454 | Bacteriophage Gp15 | 243 | 44% | 44% | 42% | 40% |
| 27470 | Phage prohead protein | 1872 | 64% | 54% | 45% | 52% |
| 29331 | Phage portal protein | 1254 | 69% | 67% | 60% | 60% |
| 30695 | Terminase | 1800 | 62% | 60% | 58% | 58% |
